# Sexual Dimorphism and Geographic Variation in Washington Bobcat (*Lynx rufus*) Skulls

**DOI:** 10.64898/2026.09.09.750527

**Authors:** Alise Newman, Chris J. Law

**Affiliations:** Department of Biology and Burke Museum, University of Washington, Seattle, WA, USA

## Abstract

Sexual dimorphism is a well-studied phenomenon in which males and females of a species exhibit different phenotypic characteristics. However, the degree of sexual dimorphism within a species may vary among populations and may be influenced by geographic factors such as seasonality, prey availability, and population density. In this study, we examined the effects of sexual dimorphism and geographic region on the craniomandibular morphology of bobcats (*Lynx rufus*) in Washington State. Using digital calipers, we measured 13 craniomandibular traits from 35 males and 42 females. We found that the degree of skull dimorphism does not differ between the eastern and western side of the Cascade Mountains. However, we found significant effects of sex and geographic region on craniomandibular morphology. Across the geographic regions, males were significantly larger than females in most craniomandibular traits, excluding postorbital constriction breadth (POC). After correcting for expected size differences, only condylobasal length (CBL) was disproportionately longer in males, indicating subtle shape dimorphism within this trait, that is largely unexplained by allometry. Mechanical advantage (MA) of the temporalis and masseter muscles did not differ between the sexes at the canine or the molar bite points. Comparisons between populations east and west of the Cascade Range revealed that while sex remains the primary indicator of overall size, there is a small regional effect on size as western bobcats are slightly larger than eastern bobcats. These regional differences highlight how environmental pressures including local prey availability and severe winter conditions may influence functional skull design. Future ecological and behavioral studies are needed to further clarify how these localized environmental drivers shape phenotypic variation across populations.

## Introduction

Sexual dimorphism (SD), or phenotypic differences between males and females of a species, is widespread among mammals. SD has been intensively studied in carnivorans, revealing that males tend to have larger, heavier bodies and often craniomandibular traits that enhance biting ability compared to females (Lindenfors et al. 2007; Morris and Carrier 2016; Christiansen and Harris 2012). Bobcats (*Lynx rufus)* are no exception, exhibiting male-biased size dimorphism. Bobcats are medium-sized felids that can subdue prey exceeding its size (Segura 2015). Bobcats have a widespread distribution across North America and can survive in a variety of habitats (Larivière and Walton 1997), making them an ideal model species to study regional and climatic influences on craniomandibular morphology. Characterized by its short tail and tawny brown coat with black spots, they rely heavily on effective camouflage through vegetated cover to stalk and hunt prey, often avoiding areas of intense agricultural development (Hansen 2006). Male bobcats can be 25-80% heavier (Larivière and Walton 1997) and 6-9% larger than females in linear measurements of the skull (Dobson and Wigginton 1996). While SD is evident in the bobcat skull, the mandible often contains a greater degree of dimorphism than that of dental or cranial measurements (Albert 1981). As with most solitary, obligate carnivorans, SD is generally hypothesized to result from sexual selection, specifically competition between males for territories and access to females under a polygynous mating system (Darwin 1871; Clutton-Brock 2007). Alternatively, the niche divergence hypothesis suggests that intersexual competition for limited resources may contribute to the evolution or maintenance of morphological dimorphism alongside resource partitioning between the sexes (Shine 1989; Selander 1966; Hedrick and Temeles 1989; Law and Mehta 2018). Finally, macroecological factors such as seasonality, latitude, population density, or topography may play a role altering magnitudes of sexual dimorphism as selection on body size may impact the sexes to different degrees (Dobson and Wigginton 1996; Isaac 2005; Sikes and Kennedy 1993).

SD is well demonstrated in bobcats across the United States, but minimal research has been conducted to explain differences in SD between vastly different geographic regions. Thus, in this study, we examine differences in craniomandibular traits between male and female bobcats across Washington State and investigate how niche divergence and seasonality may influence sexual dimorphism. Washington provides a great model system because it encompasses two distinct geographic cohorts of bobcats separated by the barrier of the Cascade Mountain crest: the coastal *Lynx rufus fasciatus* to the west (hereinafter referred as western bobcats) and *Lynx rufus pallescens* to the east (hereinafter referred as eastern bobcats) (Hall and Kelson 1959; Knick et al. 1985; Hansen 2006). Historically, these populations have been treated as distinct subspecies with differences in diet, habitat type, and degree of sexual dimorphism.

Washington exhibits strong east-west differences in seasonality across the Cascade Range, with greater seasonality and colder winter temperatures in eastern Washington as a result of the rain shadow effect (Mass 2021; Parson et al. 2001). Modern range-wide taxonomic revisions have unified these groups under a single western subspecies designation (*L. r. fasciatus*) (Kitchener et al. 2017), as the Cascade Crest has been shown not to be an absolute barrier to gene flow in bobcats (Reding et al. 2013). Despite this, the stark differences in availability of resources and climatic divide across the Cascades may result in morphological variation between the two geographic regions.

The niche divergence hypothesis presents a possible explanation for differences in size dimorphism between the sexes. Bobcats exhibit differences in prey selection between the sexes, in which females take smaller prey items such as lagomorphs, whereas males more frequently take larger prey items such as deer (Litvaitis et al. 1984; Baker et al. 2001). In environments with limited resources, competition between males and females may drive natural selection resulting in a differential prey selection (McLean et al. 2005) and morphological differences in the feeding apparatus. As competition is generally greater for vertebrate prey than nonvertebrate prey, size dimorphism is generally greater in solitary, carnivorous species (Law 2019). Bobcats are known to have partial to complete overlap of male-female home ranges and therefore dietary differences may reduce intraspecific competition (Litvaitis et al. 1984). In western Washington, a bobcat’s diet primarily consists of mountain beavers and snowshoe hares, whereas the eastern population has a more variable diet, consisting of lagomorphs, red squirrels, deer and voles (Knick et al. 1984). Additionally, eastern Washington bobcats have a larger niche breadth than western Washington bobcats, indicating a more generalized and variable diet (Newbury and Hodges 2018). This greater diversity in prey selection may be due to food-limitations in the eastern populations compared to the western populations (Knick et al. 1984). Bobcats of eastern Washington exhibit larger home ranges and lower population densities than their western counterparts (WDFW 2026), likely due to reduced prey availability (McLean et al. 2005; Knick et al. 1985; Larivière and Walton 1997). In turn, less prey availability may contribute to increased competition between the sexes in eastern Washington, which may result in higher levels of SD (Li and Kokko 2021).

Geographic variation, specifically seasonality, may also influence sexual dimorphism in carnivorans. Across 101 terrestrial mammalian carnivores, primary productivity and environmental seasonality were determined to be primary drivers of body size and sexual dimorphism; however, these climatic forces do not act directly on morphology but are instead a result of changes in spacing behavior such as home range size and population density (Ferguson and Larivière 2008). Environmental seasonality can alter magnitudes of sexual dimorphism through two possible pathways. First, greater seasonality could increase dimorphism through a combination of Bergmann’s rule, which predicts that body size in all individuals is larger in colder climates (Bergmann 1847), and Rensch’s rule, which states the magnitude of SD increases with increasing body sizes (Rensch 1950). Thus, when these two rules intersect, individuals or populations occupying colder, more seasonal, regions may exhibit higher degrees of SD (Blanckenhorn et al. 2006, Ferguson and Larivière 2008). While Bergmann’s rule traditionally associates larger body sizes with high latitudes in colder temperatures (Bergmann 1847), patterns in body size may be more closely driven by the degree of seasonality and resource predictability (Quin et al. 1996). Bobcats across North America have been shown to follow Bergmann’s rule (Wigginton and Dobson 1999), as do others within the family Felidae such as the mountain lion (*Puma concolor*) (Kurtén 1973). It is unknown whether this trend appears across differing climatic ecoregions within Washington State. Conversely, prior research has shown that bobcats in regions of greater seasonality display lower levels of sexual dimorphism (Dobson and Wigginton 1996). A possible explanation is that regions with highly seasonal climates and deep snow accumulation often exhibit decreased prey availability, making it is more difficult for bobcats to capture prey due to their lack of long legs and snowshoe-like feet, found Canadian lynx (McLean et al. 2005, Applegate and Bahrt 1993). As a result, highly seasonal climates may minimize the degree of sexual dimorphism due to selection for larger body sizes, which may allow a species to endure longer periods of fasting (Dobson and Wigginton 1996, Millar and Hickling 1990). As males are the larger sex, energetic constraints on males may limit their body size, thereby placing an upper bound on male size (Blanckenhorn 2000) while females must maintain a minimum size to survive winter fasting (Dobson and Wigginton 1996), thus minimizing dimorphism between the sexes. This trend has been shown to occur in other mammals such as the red deer (*Cervus elaphus*), in which a colder climate was associated with decreased dimorphism (Post et al. 1999).

Here, we examined the effects of sexual dimorphism and geographic variation on bobcat craniomandibular morphology and the mechanical advantage of bite force across Washington State. We first tested the hypothesis that the degree of craniomandibular SD differs between the eastern and western bobcats. We predict that craniomandibular SD will be greater in eastern bobcats than western bobcats because greater resource competition often correlates with higher levels of sexual dimorphism (Li and Kokko 2021). Alternatively, if seasonality is the primary driver of morphology we predict one of two patterns: 1) eastern bobcats will exhibit greater SD if increased body size from Bergmann’s rule interacts with Rensch’s rule in highly seasonal climates such as in eastern Washington (Blanckenhorn et al. 2006) or 2) eastern bobcats will exhibit reduced SD if winter energetic constraints and fasting demands select for larger body sizes, while imposing an upper limit on male body size and lower limit on female body size (Dobson and Wigginton 1996). To test these predictions, we measured 13 craniomandibular traits from 35 males and 42 females across eastern and western Washington. We then investigated how sexual dimorphism and geographical regions affected variation in these traits. We also tested differences in mechanical advantage between males and females to explore if differential prey selection may be associated with morphology, specifically the feeding apparatus.

## Methods

We obtained 77 bobcat (*Lynx rufus*) skulls (35 males and 42 females) from the Burke Museum of Natural History and Culture, the Puget Sound Museum of Natural History, and the Charles R. Conner Museum. All specimens were collected from 16 localities across Washington State spanning regions east and west of the Cascade Range. Of the 77 specimens, we were able to sample 48 bobcats from the western side of the Cascade Range (historically *L. r. fasciatus*) and 29 bobcats from the eastern side of the Cascade Range (historically *L. r. pallescens*). Only adult specimens were used in this study, determined by the fusion of cranial sutures such as the basioccipital-basisphenoid suture. Using digital calipers, we measured eight cranial and five mandibular traits (Figure 1; Table 1) to identify size differences between male and female specimens. We also estimated the overall size of the skull by calculating the geometric mean size (hereinafter called skull size) for each individual using all 13 traits. The geometric mean is widely used as a proxy for size and is derived from the 13th root of the product of the 13 linear measurements (Mosimann 1970; Jungers et al. 1995). All morphometric data processing, statistical testing, and visualizations were performed in R (version 4.4.2) within the RStudio integrated development environment (version 2024.12.0+467). All raw dataset files and analysis code scripts are provided as Supporting Information (S1 File and S2 File).

**Figure 1.**
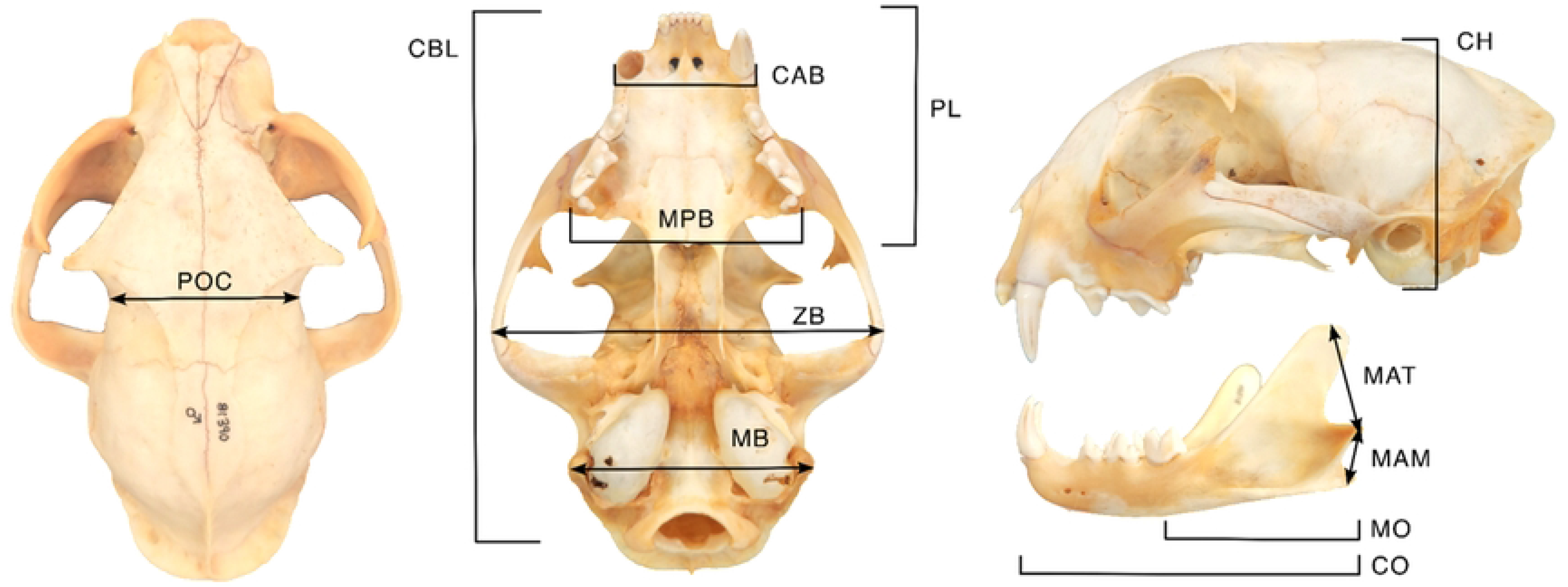
Measurements of the 13 craniomandibular traits used for evaluation of sexual dimorphism in bobcats (*Lynx rufus*).

**Table 1.**
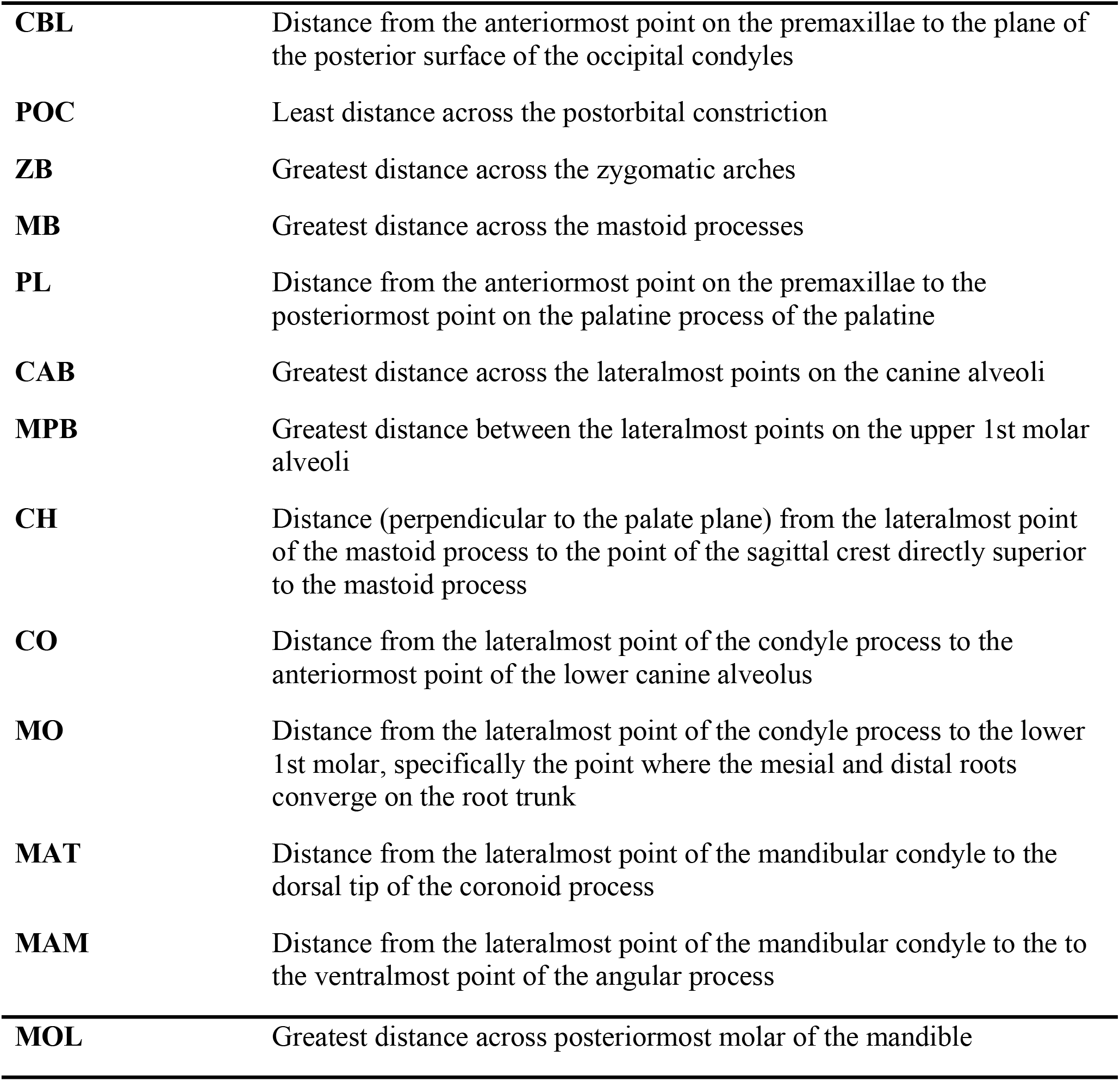
Craniomandibular measurements used to evaluate sexual dimorphism in bobcats *(Lynx rufus).* CBL = condylobasal length; POC = postorbital constriction breadth; ZB = zygomatic breadth; MB = mastoid breadth; PL = palatal length; CAB = canine alveoli breadth; MPB = maximum palatal breadth; CH = cranial height; CO = canine out-lever; MO = molar out-lever; MAT = moment arm of temporalis (in-lever); MAM = moment arm of masseter (in-lever); MOL = molar.

### Analysis of sexual dimorphism and geographic region on skull size and morphology

We first tested the effects of sexual dimorphism and geographic region on skull size using a two-way analysis of variance (ANOVA). Similarly, we tested the effects of sexual dimorphism and geographic region on all 13 traits using a two-way multivariate analysis of variance (MANOVA) with Pillai’s trace as a test statistic. For both models, we included an interaction between sex and geographic region to determine whether the magnitude of SD in skull size and craniomandibular morphology between males and females differs between regions. A significant interaction would indicate that SD varies geographically. We then performed a linear discriminant analysis (LDA) with leave-one-out cross-validation to determine how well the 13 traits can be predicted between the sexes and two geographic regions. The trait dataset was natural logged transformed prior to running the statistical tests.

### Analysis of size-corrected sexual dimorphism and geographic region on skull size and morphology

Because size dimorphism is prevalent in bobcats, we repeated the above analyses (i.e., two-way MANOVA and LDA) using size-corrected craniomandibular traits. To correct for size, we calculated residuals of each trait from a linear model between each trait and skull size, which represent the relative size of the trait independent of overall size. In this context, size is defined as the variation produced by allometry, and shape as the size-corrected morphological variation unexplained by allometry.

### Analysis of sexual dimorphism in individual traits

Additionally, we broke down the analysis trait by trait, conducting individual two-way ANOVAs with sex and geographic region as main effects for both the raw, log-transformed traits and the size-corrected traits. We quantified the degree of sexual dimorphism in each craniomandibular trait by calculating the SDI value based on Lovich and Gibbon’s (1992) size dimorphism index:

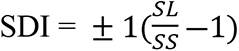

where SL and SS are the sizes of the larger and smaller trait, respectively. A negative sign is given if the male trait is larger and a positive sign given if the female trait is larger. Higher values of SDI indicate greater dimorphism and SDI values approaching zero indicate minimal sexual differences.

### Mechanical Advantage

We assessed jaw closure between the sexes by calculating the mechanical advantage (MA) of the temporalis and masseter muscles at two bite points, the canine and the first lower molar (carnassial). Mechanical advantage models the lower jaw as a lever and measures the amount of jaw force transmitted to the bite point, which can be used as a proxy for bite force. Each bite point serves a different functional behavior, with the canine designed for prey capture and the molars designed for prey processing and chewing (Salcido and Polly 2026). While males may require greater bite force at the canines to obtain larger prey, both sexes need to process prey efficiently at the molars, which may be a factor influencing varying degrees of sexual dimorphism at different parts of the mandible. To calculate the MA of the temporalis (MA_tem_), we measured the ratio between the temporalis in-lever (the distance between the mandibular condyle and the dorsal tip of the coronoid process) and the out-lever (the distance between the mandibular condyle and the bite point). To calculate the MA of the masseter (MA_mas_), we measured the ratio between the masseter in-lever (the distance between the mandibular condyle and the ventralmost point of the angular process) and the out-lever. We used the lower first molar and the canine as bite points to measure MA, as they are necessary for carnivorans to grind and slice prey. Two-way ANOVA tests were used to determine statistical differences in the mechanical advantage between the sexes and between the two regions.

## Results

### Differences in skull size between regions and sexes

We did not find a significant interaction between sex and geographic region in skull size (Table 2, A), indicating that the overall degree of sexual dimorphism in skull size does not differ east and west of the Cascade Range. Both sex and geographic region exhibited significant effects on skull size (Table 2, A), where males (42.24 mm) were significantly larger than females (38.91 mm) and western bobcats (40.64 mm) were significantly larger than eastern bobcats (40.07 mm) (Figure 2; Table 2, A). Skull size in males was approximately 8.3% larger than females (coefficient estimate =0.083, SE = 0.007). Skull size in western bobcats was approximately 1.8% larger than eastern bobcats (coefficient estimate =0.018, SE = 0.007).

**Figure 2.**
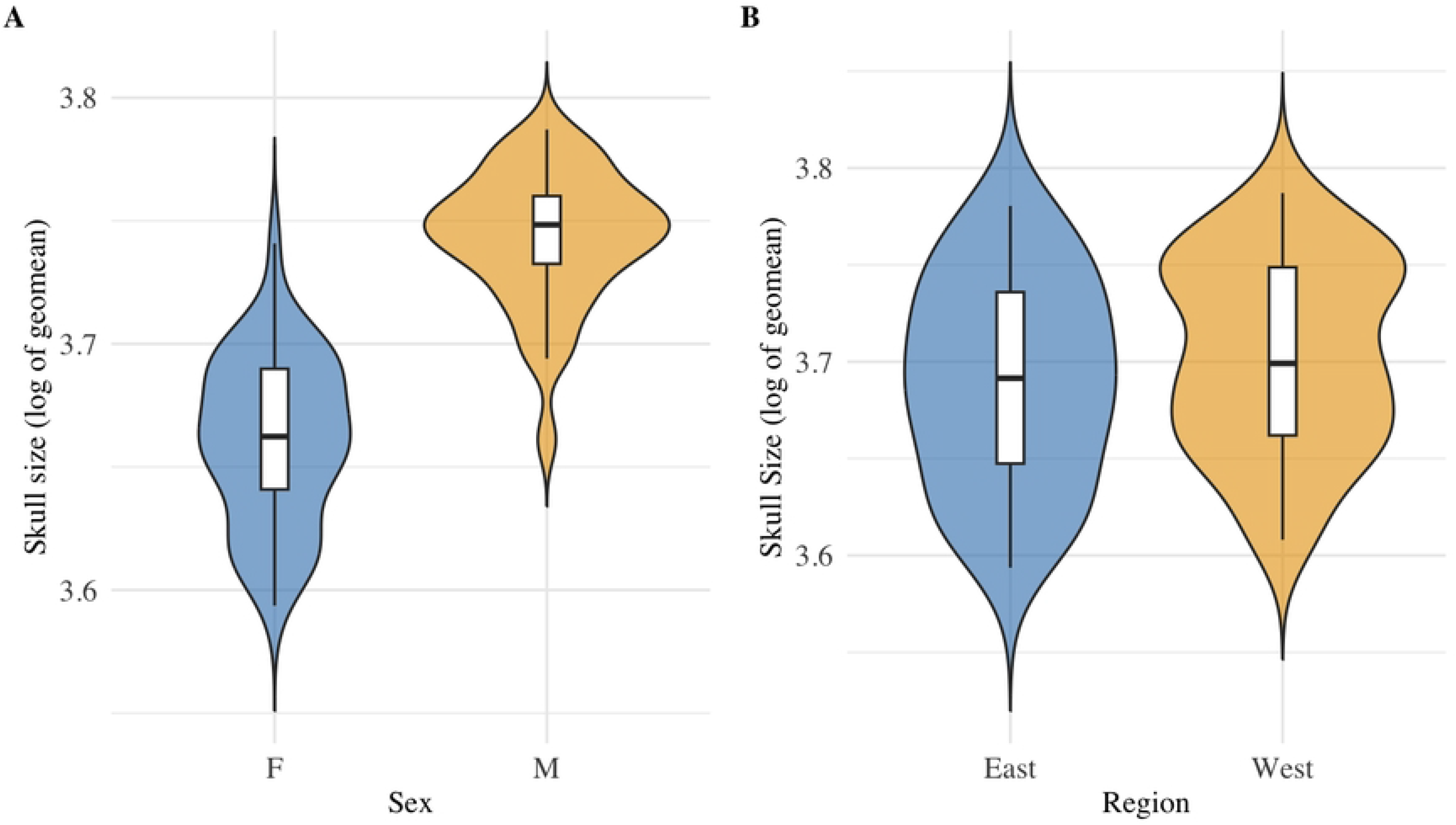
Regional and sex-specific variation in bobcat *(Lynx rufus)* skull size in Washington State. (A) Sexual size dimorphism showing log-transformed skull size between females (n = 42) and males (n= 35). (B) Comparison of log-transformed skull size between eastern (n=29) and western (n=48) Washington populations.

**Table 2.** ANOVA and MANOVA statistics for skull size and overall craniomandibular traits using raw, log-transformed traits (A, B) and size-corrected traits (C). Bold P-values indicate significance (P < 0.05).

| A) SD in skull size (two-way ANOVA) | F<sub>1,73</sub> | Pvalue |
| --- | --- | --- |
| Sex | 148.63 | **P < 0.001** |
| Georegion | 6.57 | **P = 0.012** |
| Interaction (Sex*Georegion) | 0.509 | P = 0.478 |
| **B) SD in craniomandibular morphology (two-way MANOVA)** | **F<sub>13,61</sub>** | **Pvalue** |
| Sex | 16.41 | **P < 0.001** |
| Georegion | 3.62 | **P < 0.001** |
| Interaction (Sex*Georegion) | 0.642 | P = 0.809 |
| **C) Size-corrected SD in craniomandibular morphology (two-way MANOVA)** | **F<sub>12,62</sub>** | **Pvalue** |
| Sex | 0.922 | P = 0.531 |
| Georegion | 3.93 | **P = 0.002** |
| Interaction (Sex*Georegion) | 0.671 | P = 0.772 |

### Differences in craniomandibular morphology between regions and sexes

Similarly, we did not find a significant interaction between sex and geographic region in overall craniomandibular morphology (Table 2, B). Both sex and geographic region exhibited significant effects on overall craniomandibular morphology (Table 2, B). The LDA with cross-validation reclassified 64.9% of individuals in the correct sex*geographic region category (Table 3, A; Figure 3, A). Percentages of correctly reclassified sex and geographic region were 93.5% and 70.1%, respectively.

**Figure 3.**
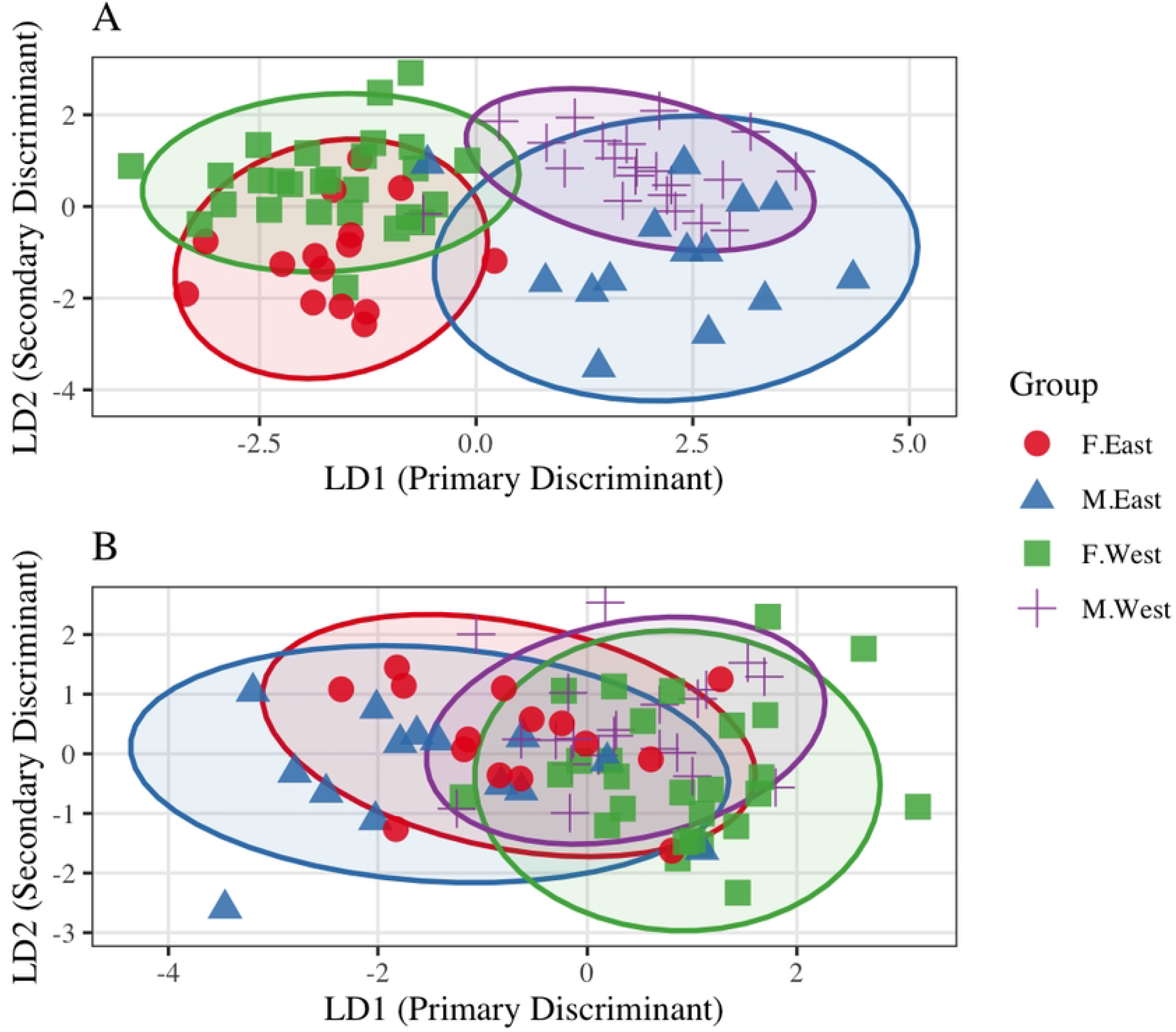
Linear Discriminant Analysis (LDA) of cranial variation across sex and region groups. (A) 13 log-transformed raw traits (Cross-validated accuracy: [64.9]%). (B) Size-corrected traits (Cross-validated accuracy: [31.2]%). Axis LD1 separates individuals by sex (female vs. male), and axis LD2 separates individuals by geographic region (east vs. west). Ellipses represent 95% data boundaries.

**Table 3.** Cross validated LDA confusion matrices (n = 77) detailing the classification accuracy of A) raw, log transformed traits (size dimorphism) and B) size-corrected traits (shape dimorphism). Rows indicate the true sex and geographic region of the individuals and columns indicate the predicted group assignment. Bold values along the diagonal indicate correct classifications. Removing the effect of overall size reduces the classification accuracy from 64.9% to 31.2%, indicating that overall size, rather than shape, is the primary driver of morphological differences between sexes and regions.

**A. Craniomandibular morphology**
| Actual sex.region | Predicted F.East | Predicted M.East | Predicted F.West | Predicted M.West |
| --- | --- | --- | --- | --- |
| F.East | **8** | 1 | 6 | 0 |
| M.East | 1 | **7** | 1 | 5 |
| F.West | 7 | 0 | **20** | 0 |
| M.West | 0 | 4 | 2 | **15** |

**B. Size-corrected craniomandibular morphology**
| Actual sex.region | Predicted F.East | Predicted M.East | Predicted F.West | Predicted M.West |
| --- | --- | --- | --- | --- |
| F.East | **2** | 6 | 3 | 4 |
| M.East | 5 | **5** | 1 | 3 |
| F.West | 4 | 1 | **15** | 7 |
| M.West | 3 | 1 | 15 | **2** |

### Analysis of size-corrected sexual dimorphism and geographic region on craniomandibular morphology

After size-correcting our trait data, we found that only geographic region exhibited a significant effect on size-corrected craniomandibular morphology (Table 2, C). We did not find a significant interaction between sex and geographic region or an effect of sex on size-corrected craniomandibular morphology (Table 2, C). To prevent matrix singularity caused by the collinearity and zero-sum constraint of the geometric-mean calculation, we dropped a single redundant trait before running the Type II MANOVA using the size-corrected traits. The LDA using size-corrected traits reclassified 31.17% of individuals in the correct sex*geographic region category (Table 3, B; Figure 3, B). Percentages of correctly reclassified sex and geographic region were 45.5% and 74.0%, respectively.

### Analysis of skull sexual dimorphism in individual traits

Breaking down the analysis trait by trait, individual ANOVA tests revealed all craniomandibular traits, except for postorbital constriction breadth, were significantly larger in males than females (Table 4). The CBL showed to have the largest standard deviation among the specimens. SDI values ranged from -2.0% (POC) to -12.7% (MAT). The palatine length (PL), canine alveoli breadth (CAB), molar out-lever (MO), and size of posteriormost molar (MOL) were found to have significant regional differences with all traits larger in the western population.

**Table 4.** Descriptive statistics of craniomandibular traits of male and female bobcats (*Lynx rufus*) prior to size correction. Trait values are in millimeters. Fstat and Pvalues from ANOVA to test for differences in means for each measurement between the sexes. Bold P-values indicate significance (P < 0.05). CBL = condylobasal length; POC = postorbital constriction breadth; ZB = zygomatic breadth; MB = mastoid breadth; PL = palatal length; CAB = canine alveoli breadth; MPB = maximum palatal breadth; CH = cranial height; CO = canine out-lever; MO = molar out-lever; MAT = moment arm of temporalis (in-lever); MAM = moment arm of masseter (in-lever); MOL = molar.

| Trait | Sex | Mean | SD | Min | Max | F<sub>1,73</sub> | Pvalue | SDI |
| --- | --- | --- | --- | --- | --- | --- | --- | --- |
| GMS | F | 38.909 | 1.304 | 36.368 | 42.131 | 148.63 | <0.0001 | -0.086 |
|  | M | 42.245 | 1.133 | 38.904 | 44.128 |  |  |  |
| CBL | F | 108.902 | 3.265 | 101.700 | 113.930 | 191.69 | <0.0001 | -0.100 |
|  | M | 119.798 | 3.498 | 111.150 | 127.100 |  |  |  |
| ZB | F | 85.538 | 2.805 | 79.640 | 91.790 | 144.78 | <0.0001 | -0.095 |
|  | M | 93.638 | 3.109 | 86.550 | 101.270 |  |  |  |
| POC | F | 39.889 | 1.690 | 34.330 | 43.210 | 3.29 | 0.074 | -0.020 |
|  | M | 40.669 | 1.916 | 36.400 | 43.680 |  |  |  |
| MB | F | 53.856 | 1.426 | 50.680 | 56.480 | 114.72 | <0.0001 | -0.069 |
|  | M | 57.561 | 1.551 | 53.270 | 60.240 |  |  |  |
| PL | F | 46.432 | 1.894 | 40.280 | 50.130 | 121.47 | <0.0001 | -0.096 |
|  | M | 50.880 | 1.745 | 47.700 | 54.600 |  |  |  |
| CAB | F | 31.313 | 1.877 | 27.800 | 35.520 | 61.62 | <0.0001 | -0.092 |
|  | M | 34.202 | 1.817 | 30.100 | 37.360 |  |  |  |
| MPB | F | 47.405 | 2.004 | 43.640 | 53.000 | 66.55 | <0.0001 | -0.078 |
|  | M | 51.092 | 1.937 | 47.050 | 55.600 |  |  |  |
| CH | F | 43.371 | 0.966 | 41.640 | 45.280 | 50.55 | <0.0001 | -0.043 |
|  | M | 45.239 | 1.319 | 43.100 | 48.270 |  |  |  |
| CO | F | 75.147 | 2.499 | 70.420 | 80.070 | 169.39 | <0.0001 | -0.105 |
|  | M | 83.056 | 2.872 | 75.020 | 87.960 |  |  |  |
| MO | F | 40.862 | 1.881 | 36.010 | 44.580 | 97.19 | <0.0001 | -0.110 |
|  | M | 45.340 | 2.155 | 39.600 | 49.330 |  |  |  |
| MAT | F | 22.937 | 1.349 | 20.340 | 26.170 | 69.66 | <0.0001 | -0.127 |
|  | M | 25.841 | 1.691 | 22.180 | 29.850 |  |  |  |
| MAM | F | 10.445 | 1.175 | 8.120 | 12.750 | 19.61 | <0.0001 | -0.105 |
|  | M | 11.543 | 1.082 | 9.500 | 14.460 |  |  |  |
| MOL | F | 10.756 | 0.719 | 9.490 | 12.570 | 33.73 | <0.0001 | -0.081 |
|  | M | 11.623 | 0.668 | 10.320 | 13.370 |  |  |  |

When examining the relative size of each trait (i.e., size-corrected traits), we found no significant sexual dimorphism across the skull (Table 2, C), indicating a lack of shape dimorphism. Individual ANOVAs further revealed that only the CBL remained significantly larger in males after size correction (P = 0.014). However, the effect size for this relationship was low, with sex accounting for 16.6% of the variance in residual skull length (Adjusted R^2^=0.166). After controlling for skull size, the model estimates that male CBL exceeds that of females by a value of approximately 0.76% (coefficient estimate =0.0070, SE=0.004). After size correction, three traits displayed regional differences in relative size, or changes in trait proportions. The condylobasal length (CBL) and zygomatic breadth (ZB) were relatively larger in the east, while the canine alveoli breadth (CAB) was relatively wider in the west.

### Mechanical Advantage

We measured MA using the raw, linear traits to estimate feeding performance at the two primary jaw adductor muscles, the temporalis and the masseter muscle. At both the canine and the molar bite point, mechanical advantage of either the masseter or temporalis muscle was not significant (Table 5), which indicates males and females produce an approximately uniform mechanical advantage. To visualize the variation and density distributions of jaw leverage across individuals, we mapped the raw mechanical advantage data across all four muscle and bite point combinations using a series of violin plots, which showed that mechanical advantage remains uniform across sexes (Figure 4). Across both sexes, the temporalis muscle exhibited higher raw mechanical advantage values compared to the masseter muscle at both functional bite points. The two-way ANOVA conducted at each bite point displayed no regional differences in MA, nor interaction between sex and region (Table 5).

**Figure 4.**
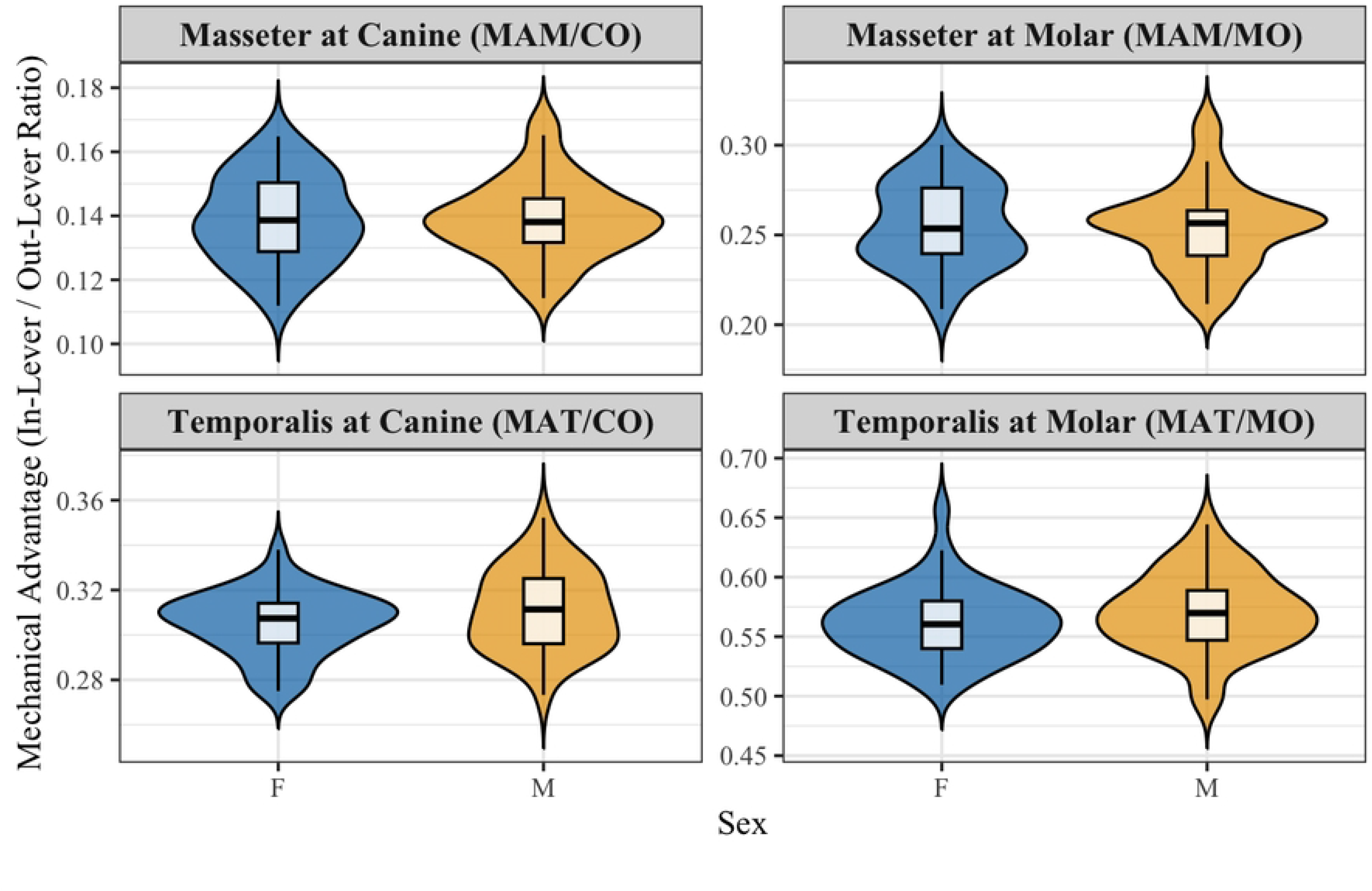
Sex-specific variation in mechanical advantage of the masseter and temporalis muscles at the canine and molar bite points.

**Table 5.**
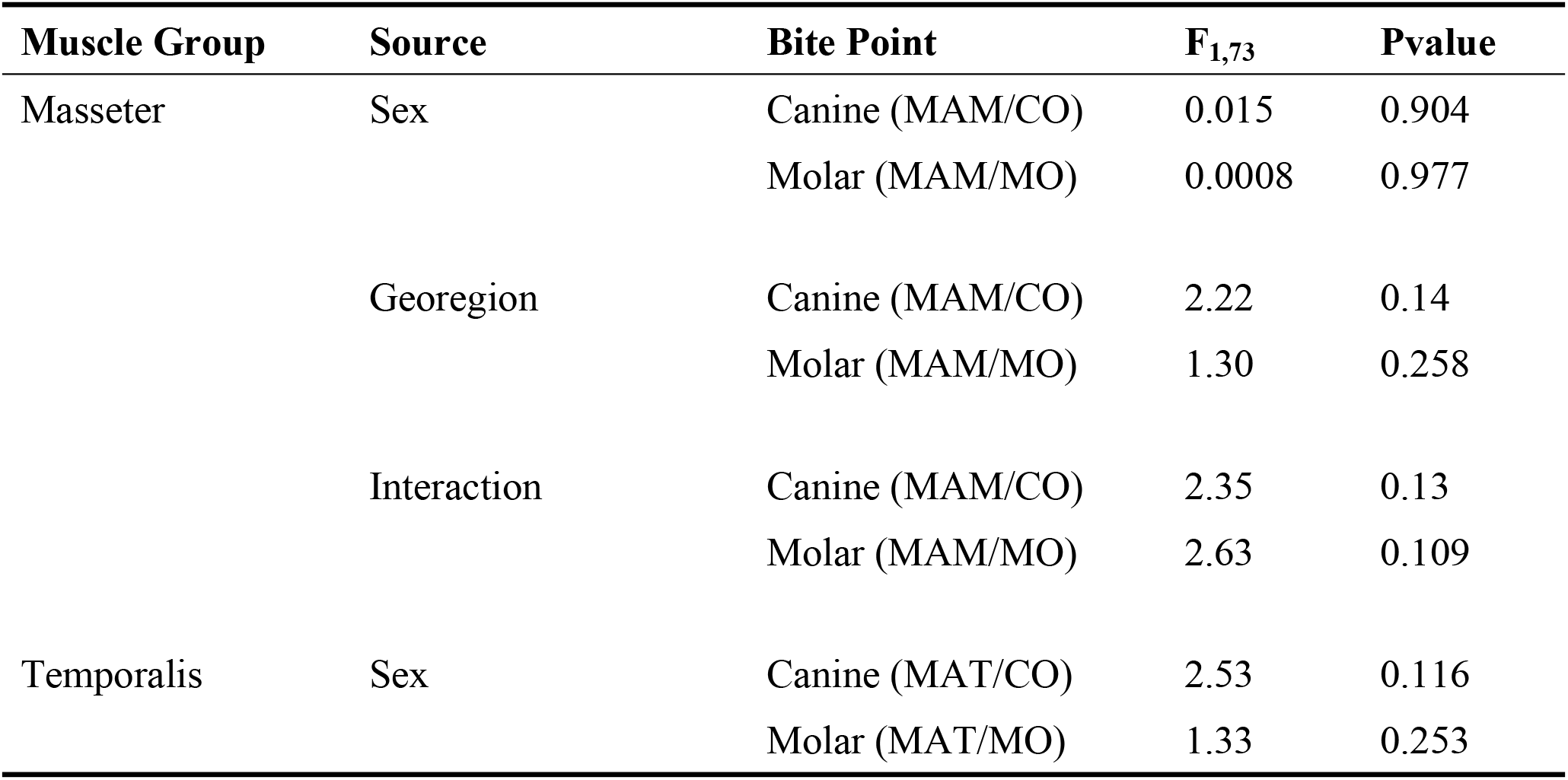
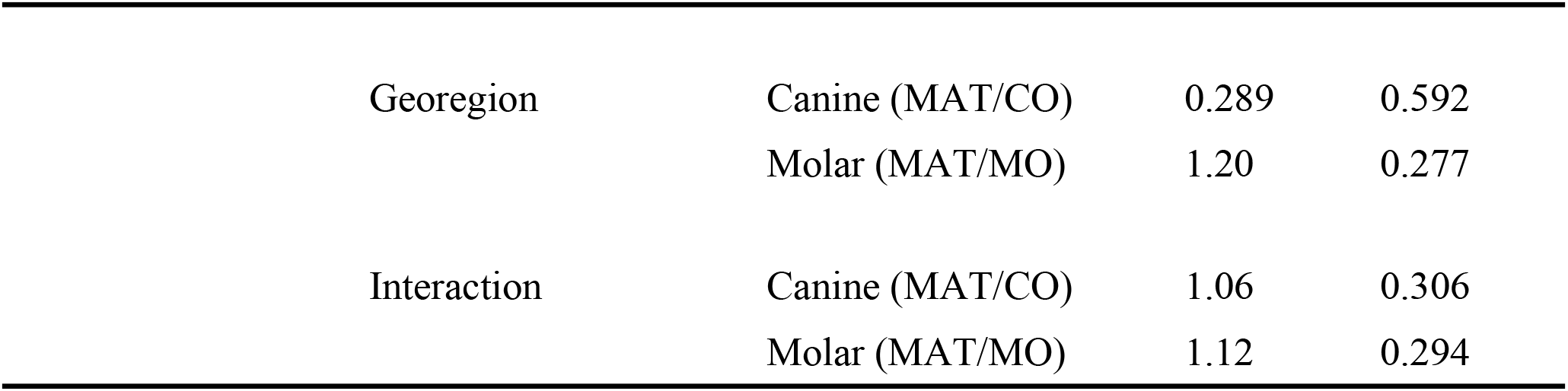
Two-way ANOVA statistics (Fstat and Pvalue) for mechanical advantage for both muscle groups across sources, and bite points. Bold P-values indicate significance (P < 0.05).

## Discussion

We found that both sex and geographic region had significant effects on skull size and craniomandibular morphology of bobcats in Washington State. However, our results do not show a statistically significant interaction effect, thus providing no evidence that craniomandibular dimorphism differs within populations east and west of the Cascade Crest.

### Assessment of the niche divergence hypothesis

We did not find support for the niche divergence hypothesis, which predicted greater magnitudes of SD in eastern Washington where food is limited, and competition is greater. Specifically, we did not find evidence that sexual dimorphism in craniomandibular morphology differs between the western and eastern regions of Washington. Furthermore, we found no evidence of changes in the magnitude of sexual dimorphism as a consequence of intersecting Bergmann’s and Rensch’s rules, which suggest populations occupying colder regions of a species distribution (such as in eastern Washington) will be larger than populations of warmer regions (Bergmann’s rule), resulting in higher degrees of SD (Rensch’s rule). Therefore, climatic differences do not create disproportionate growth rates of either sex, and the magnitude of SD remains fixed between populations.

This finding may be a result of environmental factors such as seasonality, primary productivity, and prey availability, which are likely not mutually exclusive and may drive sexual dimorphism in opposing directions. For example, under the niche divergence hypothesis, lack of food availability in eastern Washington might favor size divergence between males and females to avoid competing for the same prey, therefore increasing sexual dimorphism (Shine 1989; Selander 1966; Hedrick and Temeles 1989; Law and Mehta 2018). At the same time, Bergmann’s rule predicts that severe winter environments might select for larger-bodied individuals with smaller surface area-to-volume ratios that have a greater ability to retain body heat (Bergmann 1847). However, there is a maximum body size that can be sustained in any given environment and the steep energetic costs of maintaining large body mass could place an upper cap on male size (Huston and Wolverton 2009, Boyce 1978). Because females must maintain a minimum size to survive winter fasting (Dobson and Wigginton 1996), this upper bound on male size and lower bound on female size narrows the size gap between the sexes, thus decreasing sexual dimorphism. These competing ecological pressures ultimately may have neutralized each other to result in an equal degree of dimorphism between the two populations across Cascade Range. Further research would be necessary to systematically isolate and study the distinct effects of primary productivity, elevation, and latitude on the geographic variation of sexual dimorphism.

Before size correction, sex is the primary indicator of dimorphism in bobcat skull size, with region having a smaller effect. After size correction, sex had no effect on the shape of bobcat skulls, however, a small effect of region on the shape of the skull remains (p<0.05) exhibited in specific traits. Western bobcats exhibited larger palatine length (PL), canine alveoli breadth (CAB), molar out-lever (MO), and size of posteriormost molar (MOL). After size correction, western bobcats exhibited only a wider canine alveoli breadth (CAB). Surprisingly, the condylobasal length (CBL) and zygomatic breadth (ZB) were relatively larger in the east after size correction. While western bobcats have proportionately larger canine alveoli breadth (CAB) that may help them capture larger prey, eastern bobcats have proportionally longer skull length and width relative to their size, despite having a smaller skull size overall. While trait-by-trait variability exists between the two populations, our results revealed that the western bobcats are only 1.8% larger than eastern bobcats when considering skull size. A possible explanation for larger western bobcats may be attributed to sustained, abundant prey year-round due to the higher levels of net primary productivity on the west side of the Cascades (Law et al. 2004). It is well documented that species attain larger body sizes in areas with greater resource availability and reliable prey (Huston and Wolverton 2009, Boyce 1987). In contrast, food is scarcer in eastern Washington (Knick et al. 1984), which creates higher metabolic stress that may limit structural growth. Bobcats travel well over hard packed snow, though they lack the adaptations found in Canada lynx (*Lynx canadensis*) that allow them to efficiently hunt in deep snow, such as long legs, large snowshoe-like feet, and fur covered footpads (McLean et al. 2005, Applegate and Bahrt 1993). It may be advantageous for eastern populations to remain lighter, in order to better navigate atop snowy terrain.

### Assessment of craniomandibular morphology

We found significant evidence of sexual dimorphism in craniomandibular morphology of bobcats across Washington. Nearly all craniomandibular traits showed male-biased sized dimorphism, consistent with what has been previously found (Albert 1981). Along with body size, male-biased size dimorphism in craniomandibular morphology is considered one of the most prominent dimorphic features in carnivorans (Law et al. 2016; Gittleman and Van Valkenburgh 1997; Brunner et al. 2004; Christiansen and Harris 2012). Craniomandibular traits tend to be more robust in male carnivorans, such as total condylobasal length, size of canines, zygomatic breadth, and sagittal crest height (Gittleman and Van Valkenburgh 1997; Brunner et al. 2004; Christiansen and Harris 2012). Larger traits translate to a greater biting ability by increasing surface area for jaw muscle attachment sites (Radinsky 1981; Law 2020; Law et al. 2016). Male bobcats displaying larger craniomandibular morphology than females are consistent with these patterns across Carnivora and suggest males exhibit stronger bite force. The only trait that did not differ between the sexes was the postorbital constriction (POC) (Table 4), suggesting that either males exhibit narrower POC than expected or females exhibit wider POC than expected. Narrowing of the POC can allow for more room for the temporalis muscle, which would contribute to greater bite force (Wiig and Andersen 1986; Lynch 1996). A wider POC in females is also a possibility (Sikes and Kennedy 1993), but the benefits of that scenario may be limited. This is consistent with previous research (Albert 1981), in which postorbital breadth was identified as the sole metric lacking sexual dimorphism in size in bobcats. Across all felids, POC is identified as a constrained feature invariant between the sexes (Sicuro and Oliveira 2011).

Despite exhibiting presumably stronger bites with larger skulls, male bobcats do not exhibit significantly larger mechanical advantage (MA) at the canine or molar bite points of the masseter and temporalis muscles than female bobcats. The lack of difference between the sexes mirrors other mammalian carnivorans, such as the southern sea otter (*Enhydra lutris*), suggesting that females exhibit similar leverage efficiency of the jaw during biting despite significantly shorter out-levers (i.e., CO and MO) and jaw adductor in-levers (MAT and MAM) (Law et al. 2016), potentially due to the enhancement of different aspects of feeding mechanics resulting in similar ratios of MA. Similarly, in some rodent species such as the Talas tuco-tuco (*Ctenomys talarum*), the relationship between size and bite force is the same in both sexes, thus a higher bite force in males is a byproduct of overall size dimorphism rather than changes in jaw leverage (Becerra et al. 2011). Despite producing an approximately equal mechanical advantage, males still achieve a greater absolute bite force due to their larger cranial size and increased musculature, which allows them to consume larger prey.

In contrast, we did not find sexual dimorphism in most of the craniomandibular traits after correcting for size, suggesting that relative proportions of each trait do not differ between males and females. Significant differences in shape between the sexes were only found in CBL, in which males exhibited disproportionately longer condylobasal lengths; however, the sex effect explained only 16.6% of the variation (adjusted R^2^=0.166), indicating that this is a minor structural nuance and CBL is still mostly uniform in shape. Overall, the lack of dimorphism in these size-corrected traits suggest that male and female bobcats share similar skull shapes, and that the male skull is simply a scaled-up version of the female skull. These results are consistent the lack of mechanical advantage differences we observed. However, despite similar skull shapes and mechanical advantage, male bobcats may achieve a greater absolute bite force than female bobcats simply due to their larger size in their skull and presumably more robust jaw adductor musculature.

Why bobcats do not exhibit significant skull shape dimorphism remains unanswered as many carnivorans exhibit disproportionately bigger traits that enhance bite force such as well-developed sagittal crests, broadening of the zygomatic arches, and/or relatively larger teeth (Law 2020; Law and Mehta 2018; Gittleman and Van Valkenburgh 1997). A possible explanation is phylogenetic constraints among felids. Felids evolved relatively recently during the Late Miocene, and their hypercarnivous specializations may contribute to morphologically uniform skulls (Holliday and Steppan 2004; Segura 2015; Johnson et al. 2006). Across all modern felids, size-corrected bite force performance and mandibular mechanical advantage remains consistent irrespective of interspecific body size differences (Christiansen 2008) making the efficiency of the jaw relatively consistent (Sicuro and Oliveira 2011). The demand for a uniform, penetrating killing bite drives evolution of skull shape, which requires both a high gape and powerful bite force, which work inversely to one another, along with the enlargement of the upper canines (Christiansen 2008). Skull morphology is often divided among small felids (non-pantherine cats: lynx sp., fishing cat, and puma) and large felids (panthera cats: lion, jaguar, leopard, tiger, and snow leopard) (Werdelin 1983). While skull shape in felids is linked to absolute size and larger and smaller species show little anatomical difference, smaller felid species display larger braincase to skull volume and larger species display elongated snout regions and dorsal straightening in the posterior part of the skull, along with larger sagittal crests (Gittleman 1986; Christiansen 2008). Skull morphology within the lynx lineage consists of a broad and robust design, with some of the widest zygomatic arches and mastoid process breadths across all the felids, which maximizes space for the temporalis muscle and helps stabilize the neck during prey capture (Sicuro and Oliveira 2011). This differs from larger felids such as those in the panthera lineage, in which the snout is longer with longer rostrum, allowing a wide gape, beneficial for capturing running prey (Sicuro and Oliveira 2011).

## Conclusion

In this study we used a linear morphometric approach to examine craniomandibular dimorphism and geographical variation in bobcats across Washington State. We found no differences in the magnitude of sexual dimorphism between the two populations. Our analysis demonstrates that males are significantly larger than females in all examined metrics, excluding postorbital constriction breadth (POC), across all localities. After size correction, only condylobasal length (CBL) was disproportionately longer in males. Regionally, four traits were shown to be larger in the western population before size correction, while after size correction, two traits were relatively larger in the east and only one relatively larger in the west.

While males have a larger absolute bite force due to larger cranial size, we determined no differences in mechanical advantage between males and females, which does not provide strong support for shape-based niche divergence. Future studies should examine jaw muscle mass and overall body size scaling to determine if these factors result in dietary differences between the sexes and the two populations without altering skull lever mechanics. When comparing populations east and west of the Cascade Range, western bobcats were determined to be larger in size, which may be a result of greater year-round food availability and higher primary productivity in this region. This indicates that the ecological and selective pressures driving sexual size dimorphism, including sexual selection, resource availability, and seasonality, are not mutually exclusive, but rather may operate as counteracting forces that conserve the degree of sexual dimorphism across variable landscapes. Further studies from a wider range of regional locations across Washington would be necessary to isolate the effect of environmental variables such as seasonality and fasting endurance on skull dimorphism and overall body size.

## Acknowledgements

We are grateful to Jeff Bradley of the Burke Museum of Natural History and Culture, Gary Shugart of the Puget Sound Museum of Natural History, and Jessica Tir of the Charles R. Conner Museum at Washington State University for letting us use or loan their specimens. We thank Rita Mehta for letting A.N. collect additional data in her lab space. Generative AI and AI assisted tools (Grammarly and Google Gemini 3.6 Flash) were used for copyediting and assisting with R statistical coding during manuscript preparation. This work was funded by the United States National Science Foundation awards DEB-2447166 and DBI-2128146 (both to CJL) and the Shawn DeCew BEACON Field Research Award (to AN).

## Notes

### Competing Interest Statement

The authors have declared no competing interest.

